# A novel approach to modeling side chain ensembles of the bifunctional spin label RX

**DOI:** 10.1101/2023.05.24.542139

**Authors:** Maxx H. Tessmer, Stefan Stoll

## Abstract

We introduce a novel approach to modeling side chain ensembles of bifunctional spin labels. This approach utilizes rotamer libraries to generate side chain conformational ensembles. Because the bifunctional label is constrained by two attachment sites, the label is split into two monofunctional rotamers which are first attached to their respective sites, then rejoined by a local optimization in dihedral space. We validate this method against a set of previously published experimental data using the bifunctional spin label, RX. This method is relatively fast and can readily be used for both experimental analysis and protein modeling, providing significant advantages over modeling bifunctional labels with molecular dynamics simulations. Use of bifunctional labels for site directed spin labeling (SDSL) electron paramagnetic resonance (EPR) spectroscopy dramatically reduces label mobility, which can significantly improve resolution of small changes in protein backbone structure and dynamics. Coupling the use of bifunctional labels with side chain modeling methods allows for improved quantitative application of experimental SDSL EPR data to protein modeling.

**Statements and Declarations:** The authors declare no competing interests.

## 1. Introduction

Electron paramagnetic resonance spectroscopy (EPR) is a powerful complementary tool for the investigation of protein structure and function. Significant advances in site-directed spin labeling (SDSL) EPR, many of which were pioneered by Wayne Hubbell’s research group, have shown that EPR can be used to probe local protein structure, dynamics, topology, and membrane interactions [1–9]. The development of pulse EPR methods like double electron–electron resonance (DEER) spectroscopy [10–13] has allowed for the direct measurements of not just distances between spin labels, but the distribution of distances in a given sample [14,15], providing powerful insight into protein conformational landscapes. Such data have proven very useful for the refinement and expansion of protein models to include conformational dynamics [16–29], protein–protein docking [30–37], and even *de novo* protein folding [38–40].

Data collected from SDSL EPR experiments on a protein is necessarily determined not only by the backbone but also by the structure and dynamics of the spin label itself. Because spin labels are often very flexible, small but significant differences in protein backbone structure can be obscured by spin label side chain flexibility [41– 48]. To overcome this limitation, significant work has been done to develop conformationally restrained spin labels [49–54]. Some of the most promising include bifunctional labels, such as RX and di-histidine Cu(II) (dHisCu) [55–58]. The restrained mobility of bifunctional labels significantly improves the structural resolution of SDSL EPR experiments. Nonetheless, the side chain structure of bifunctional labels still must be considered when interpreting or utilizing experimental data. To make full quantitative use of the experimental data for protein modeling, accurate and efficient label modeling methods must be employed.

While the modeling of monofunctional labels is well developed [32,59–70], research into bifunctional label modeling has been less active [56,71–73]. Most of the modeling work of bifunctional labels has been performed using molecular dynamics (MD) and metadynamics simulations [56,68,72,73]. While these methods have proven highly accurate, they are often difficult to use and require extensive sampling at significant time and computational cost. Furthermore, force field parameterization of new labels can be tedious and require highly specialized knowledge, particularly for radicals for which very few automated parameterization tools exist. Outside of MD-based methods, to our knowledge only one method for modeling bifunctional labels has been published to date [71]. This method is specific to di-histidine copper labels and takes advantage of the flexibility of the copper coordination site. As a result, it cannot be directly generalized for covalently linked bifunctional labels like RX [55].

Here we present a novel approach to bifunctional label modeling. This approach attaches a bifunctional rotamer library and performs local optimization of each rotamer in dihedral space. We apply this method to predict distance distributions from several previously published SDSL EPR experiments using the RX label. Our results indicate that the method produces relatively good predictions with reasonable computational performance. We also analyze the performance of this method in several different backbone structures and site separation contexts to assess the generality of its utility.

## 2. Methods

### 2.1 Bifunctional Label Modeling

To model canonical amino acid side chains and monofunctional spin labels like R1 (Fig. 1A), a progenitor structure or rotamer library is superimposed on the protein backbone site of interest by aligning the structures locally around the Cα atom. This procedure does not work for bifunctional labels like RX (Fig. 1B), since they attach to two sites. Accurately aligning one site generally leads to severe misalignment of the other (Fig. 1C), and attempting to align both simultaneously leads to misalignments of both sites (Fig. 1D), even when using progenitor models generated using a similar backbone structure and site separation context as the target site pair. Forcing alignment at both sites leads to severe distortions in the side chain geometry (Fig. 1E).

**Fig. 1.**
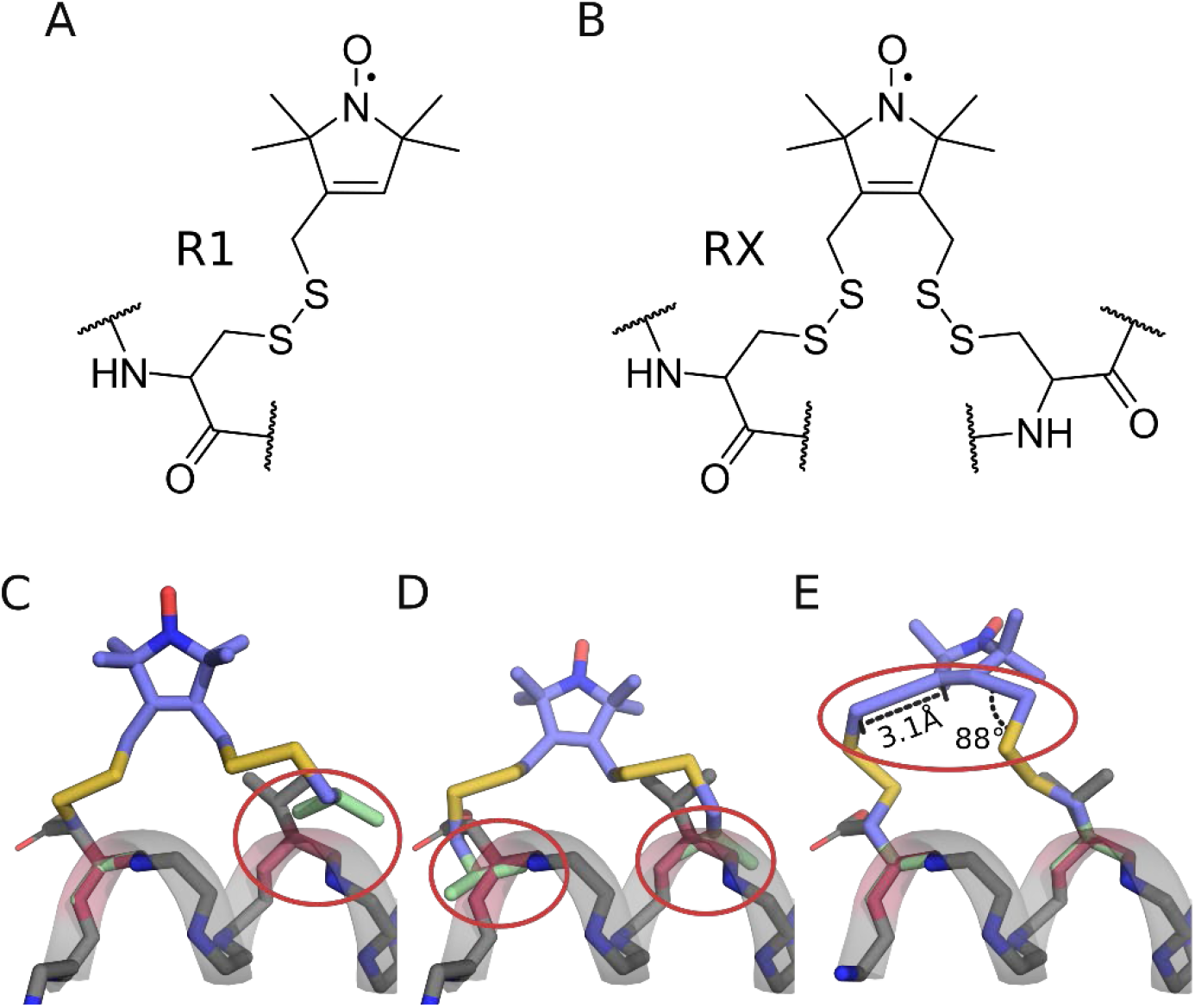
Aligning bifunctional spin labels. A) Lewis structure of the R1 spin label attached to a protein site. B) Lewis structure of the RX spin label attached to two sites. C-D) 3D structures of an RX label superimposed on an α-helix. Helix is shown as gray cartoon and sticks with gray backbone carbons. RX label is shown as sticks with light blue carbon atoms. Nitrogen, sulfur, and oxygen atoms are shown as dark blue, yellow, and light red sticks respectively. Desired backbone alignment atoms are shown in red on the helix and green on the RX label. Red circles indicate deviations in artifacts and distortions caused by the alignment method. C) 3D structure of an RX label rotamer aligned at the left site. D) 3D structure of an RX label rotamer aligned at both sites. E) 3D structure of an RX label rotamer aligned to both sites, resulting in a distorted geometry.

Our approach attempts to reconcile these unfavorable geometries by adjusting the side chain conformation via optimization of the side chain dihedral angles. Similar approaches have proven useful for the analogous problem of loop closure for protein modeling [74–76]. To achieve this, a bifunctional rotamer is represented by two separate monofunctional side chain fragments (Fig. 2B). These fragments are created by breaking the full structure of the bifunctional label at the last mobile dihedral angle of the other site. This results in two monofunctional rotamer fragments, each with a single chain of mobile dihedral angles capped with the same set of atoms after the last dihedral angle. We refer to this as the “cap”. To model the bifunctional label onto the backbone, each monofunctional rotamer fragment is locally superimposed on its target backbone site. Then, their dihedral angles are optimized to minimize the mean squared deviation between the atoms of the two copies of the cap (Fig. 2B). Breaking the side chain into two monofunctional structures and aligning the cap allows for a symmetric optimization, preventing bias of alignment errors at either attachment site. Cap misalignment is heavily penalized by a 222 kcal/mol/Å^2^force constant, chosen to be on the order of the force constant of a carbon–carbon covalent bond in the CHARMM molecular force field [77]. To prevent any internal clashes, a Lennard– Jones potential energy term is applied between all atoms of the bifunctional label that are more than 2 bonds apart (1-3 interactions). After minimization, the two monofunctional labels are merged and each cap atom is placed at the midpoint between the cap copies of the two fragments (Fig. 2B). Clashes with neighboring atoms are evaluated using a modified Lennard–Jones potential, as is commonly done when modeling monofunctional spin labels [59,78] and a small penalty is added for deviations from the starting rotamer dihedral angles at 5 kcal/mol/rad^2^. The final energy of the rotamer(s), consisting of the sum of the internal and external energies, is used to reweight the rotamer library, and low-weight rotamers are pruned from the final ensemble. If the final root mean squared deviation (RMSD) of all cap alignments in the ensemble is above 0.5 Å (0.25 Å per sub-label), we consider the modeling attempt failed. Overall, this approach utilizes dihedral flexibility in the side chain to model bifunctional labels onto site pairs, absorbing any residual misalignment into the bond stretch, angle, and dihedral angle of the last bond before the cap on both sub-labels.

**Fig. 2.**
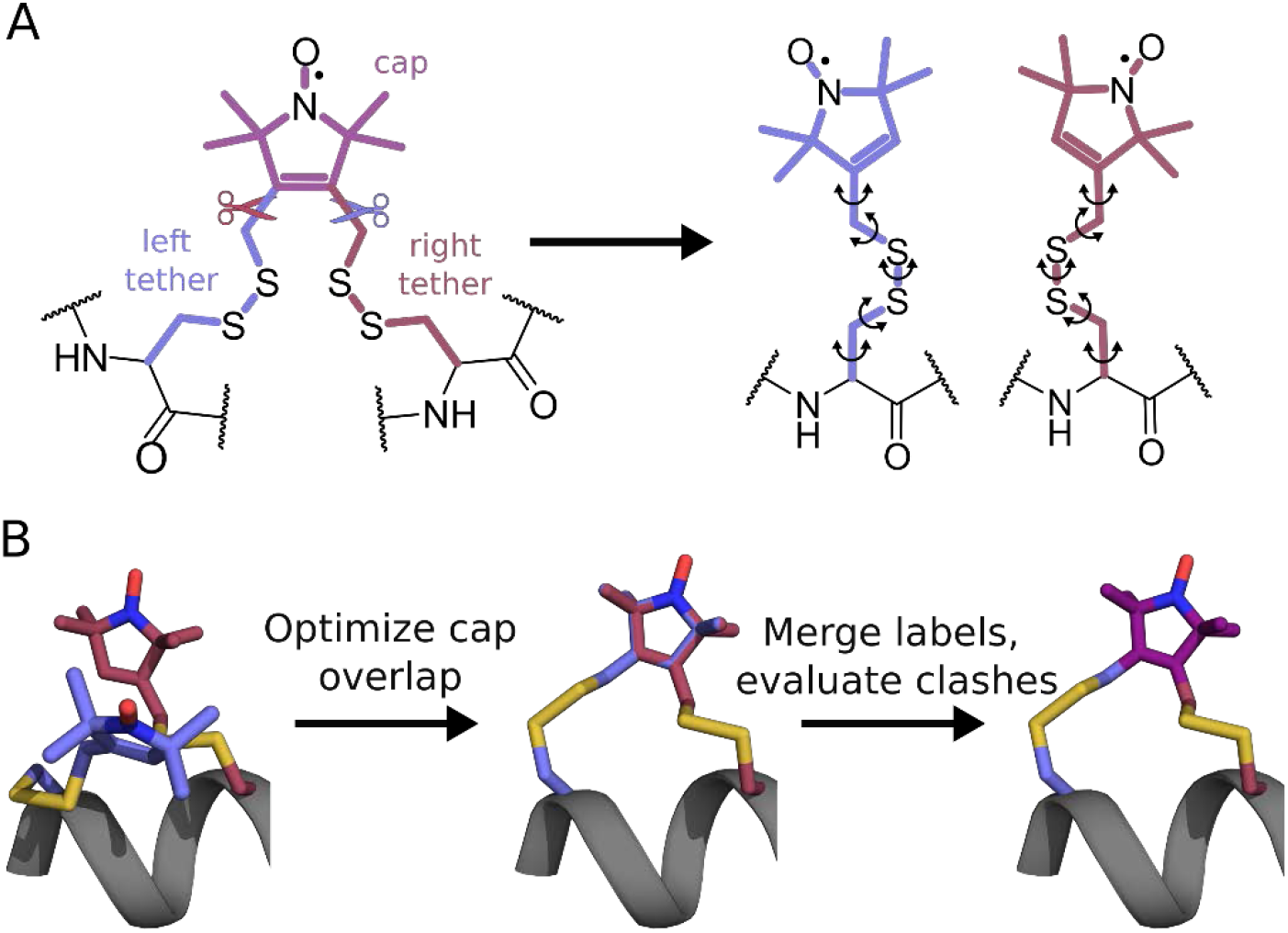
Bifunctional rotamer library modeling. A) Bifunctional labels are split into two monofunctional labels, each containing a copy of the overlapping “cap” region. B) Monofunctional labels are attached to their respective sites and the labels are optimized in dihedral space to minimize the mean squared deviation of the caps. The position of the caps are averaged and the monofunctional labels are merged into one bifunctional label. Carbon atoms of the *i*^th^ residue are shown in pale blue, and carbon atoms of the (*i*+4)^th^ residue are shown in dark red. Nitrogen, sulfur, and oxygen atoms are colored the same as in Fig 1.

Over our testing set, this process takes an average of 712 ms per RX rotamer on an Intel core i5-4670K CPU running Windows 10, This amounts to 32.8 s for the average-sized rotamer library used here (46 rotamers).

### 2.2 Bifunctional label rotamer library development

While the above methodology can be applied to a single starting structure, it is prone to get caught in a local minimum and cannot be used to generate an ensemble. Furthermore, the resulting structure does not account for stereoisomers or possible variations in bond and dihedral angles. To overcome these limitations, we utilize precalculated rotamer libraries which introduce structural variation and provide diverse starting points for minimization to better sample the conformational landscape. The utility of a precalculated rotamer library is analogous to the generation of trial conformations by *ab initio* or database methods for protein loop modeling [74–76] with the advantage of having *a priori* information on the conformational landscape of the label in a given context.

To maximize diversity in our rotamer libraries, we utilize the conformer-rotamer ensemble sampling tool (CREST) [79]. First, a PDB file is constructed to mimic the minimal system desired for modeling, in our case RX mounted onto short peptide sections. Because there are several ways of attaching the two sites of bifunctional labels like RX to a protein, we construct separate rotamer libraries for several different secondary structure and residue increment contexts, henceforth just referred to as contexts. To designate the secondary structure environment and the increment, we use the shorthand notation introduced in Fig. 3A. For example, two mounting points on the *i*^th^ and (*i*+4)^th^ residue of an α helix are referred to as an α4 context. We generated starting structures for the α3, α4, and β2 contexts by manually creating backbone templates using PyMol (version 2.4) and labeling with RX using the CHARMM-GUI input generator [72,80,81]. Each construct was capped with an N-terminal acetyl and a C-terminal amide. Glycine residues were used between the label attachment sites to minimize steric clashes with neighboring side chains. For each context, we generated rotamer libraries (Fig. 3B) using CREST [79] with the GFN forcefield [82,83] and the implicit generalized Born/surface area (GBSA) water model. Backbone torsion angles were constrained to (*φ, ψ*) = (−64°,-41°) in the α contexts and, (−135°,135°) in the β context with a cartesian force constant of 0.01 hartree/bohr^2^ (11.88 kcal/mol/Å^2^). The resulting ensembles were exported to PDB files and their relative potential energies were used to approximate populations using the Boltzmann distribution at 298 K. Since several libraries had many low population structures, often with significant redundancy, rotamers cumulatively accounting for the bottom 1% of the total population were discarded. For labeling sites whose context differs from α3, α4, and β2, we combined all context-specific libraries into a larger library, setting all rotamer weights to uniformity.

**Fig. 3.**
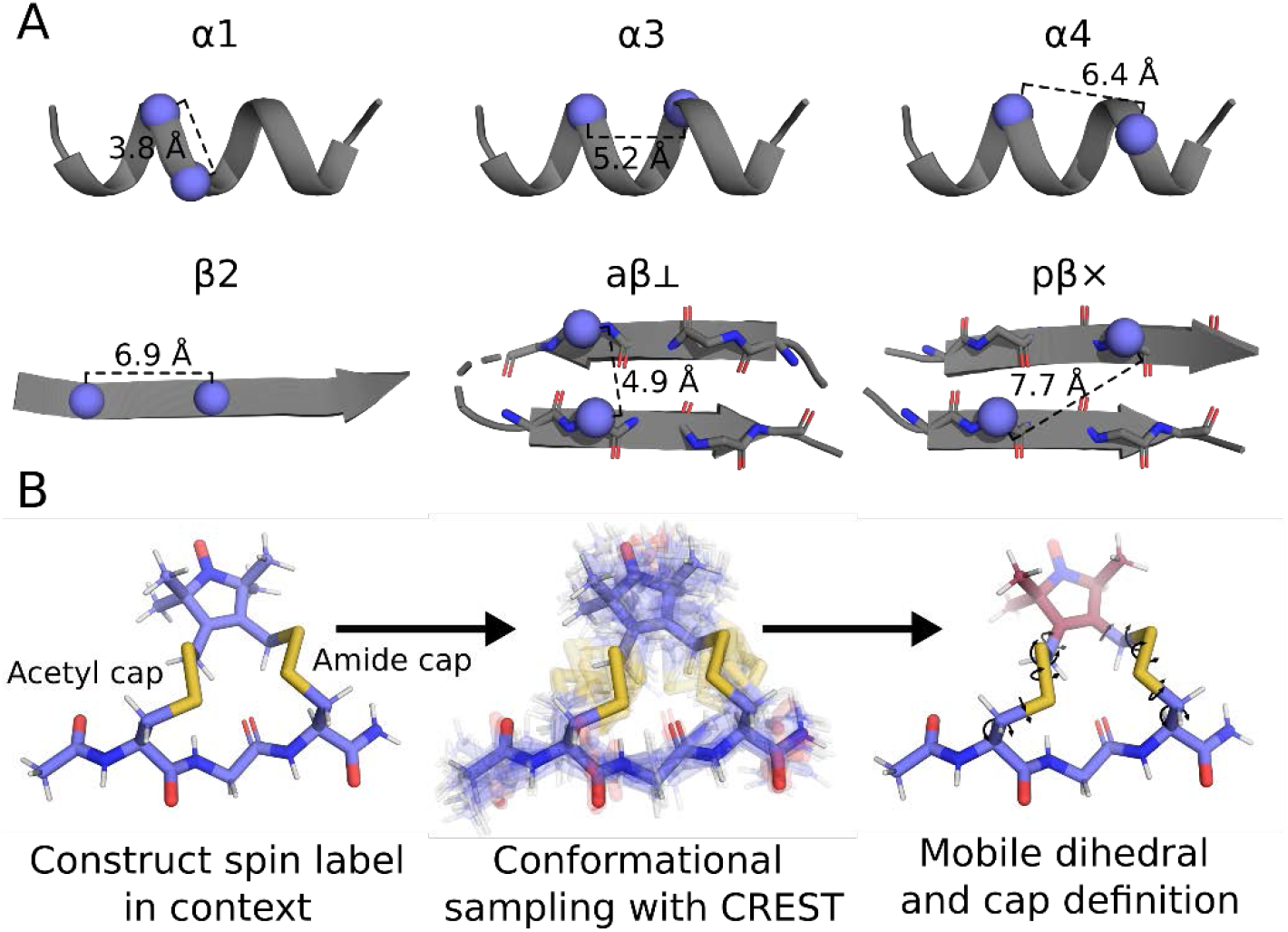
Bifunctional labeling contexts and rotamer library generation. A) Illustration of the bifunctional labeling contexts and their shorthands. “α” indicates both labeling sites are α-helical, “β” indicates both labeling sites are on β-strands or β-sheets. The number, if present, indicates the residue count between the two sites. “aβ” indicates an antiparallel beta sheet while “pβ” indicates a parallel beta sheet. ⊥ indicates that the labeling sites are perpendicular to the β-sheet, but in the same backbone hydrogen bonding register and × indicates the labeling sites are perpendicular to the β-sheet but across 2 adjacent backbone hydrogen bonding registers. B) Example of rotamer library generation process.

The bifunctional modeling method and rotamer libraries presented here are available as part of the modeling software chiLife [84] through the dRotamerEnsemble and dSpinLabel objects and can be installed from the Python package index or GitHub. Like other objects in chiLife, dRotamerEnsemble allows fine-grained control over the modeling methods, such as the minimization method, overlap penalty weights, torsion penalty weights, and more. dRotamerEnsemble and dSpinLabel objects also have many additional methods and properties that allow users to create custom analysis and protein modeling pipelines via the scriptable Python interface.

### 2.3 Analysis of DEER data

To validate the modeling approach and the rotamer libraries, we utilized experimental DEER data previously published by Stevens et al. [56] and Fleissner et al. [55]. All the raw data, generously provided by the authors, were re-analyzed using DeerLab [85]. Zero-time and phase-corrected traces were used to fit the foreground and background simultaneously with model-free Tikhonov regularization for the foreground and a homogeneous three-dimensional spin distribution model for the background. The minimum of the distance domain was 1.5 nm, and the maximum (*r*_max_) was chosen based on the maximum dipolar evolution time (*t*_max_) such that 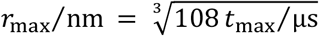 [86,87]. Modulation depth was constrained to be greater than 0 and less than 0.55. Compactness regularization was used, and the compactness and Tikhonov regularization parameters were both constrained to be between 0.01 and 10 [88,89]. These parameters were chosen based on previous results of unconstrained fitting within the lab. All fit parameters and results are reported in Figures S1-S2 and Table S1.

## 3. Results and Discussion

### 3.1 Model Predictions of DEER Experiments

To test our approach, we modeled the RX spin label on 10 site pairs with previously published experimental DEER data on the histone chaperone Vps75 [56], on T4 lysozyme (T4L) [55], and on rat intestinal fatty acid binding protein (riFABP) [55]. All the Vps75 data were intermolecular distances between singly labeled monomers under conditions that promote dimerization. Since these data were published separately, we reanalyzed all the raw data with DeerLab using a consistent protocol (Figs. S1 and S2). We used the previously published crystal structures of T4L (PDB: 2LZM) [90], riFABP (PDB: 2IFB) [91], and Vps75 dimer (PDB: 2ZD7)[92] for modeling. We divided all the DEER data into site pairs for which we had made context-specific rotamer libraries and those that used the aggregate rotamer library (see Methods). We then compared the ability of our method to predict the experimentally observed DEER distance distributions from the available crystal structures.

The results for site pairs with context-dependent rotamer libraries are shown in Fig. 4 and Fig. S3. Our method yielded good prediction accuracy, with a mean absolute displacement (MAD) of 1.3 Å between the predicted and experimental modes and a MAD of 1.7 Å between the predicted and experimental average distance. These values are consistent with modeling methods for monofunctional labels [67,82]. The largest differences between predicted and simulated distance distributions were observed in the average distance for Vps75 T122-V124RX and Vps75 A19-E23RX, deviating by 3.7 and 3.6 Å respectively. In contrast, the deviations of the mode distances are 2.1 and 0.8 Å respectively, suggesting our method predicts the predominant features of the experimental distance distributions well. In both cases, the experimental distance distributions are bimodal, skewing the average distance away from the primary mode distance. These bimodalities may arise from differences in spin label or backbone conformation. In the former case, these data suggest that the relative weights of our rotamer ensembles may not be accurately modeled. However, in the latter case, we would not expect to capture the bimodality using a single static protein model. Previous attempts to model these distributions were unable to capture the minor mode of T122-V124RX but showed some indication of modeling the minor mode of A19-E23RX [56], suggesting that both possibilities may be playing a role in our predictions.

**Fig. 4.**
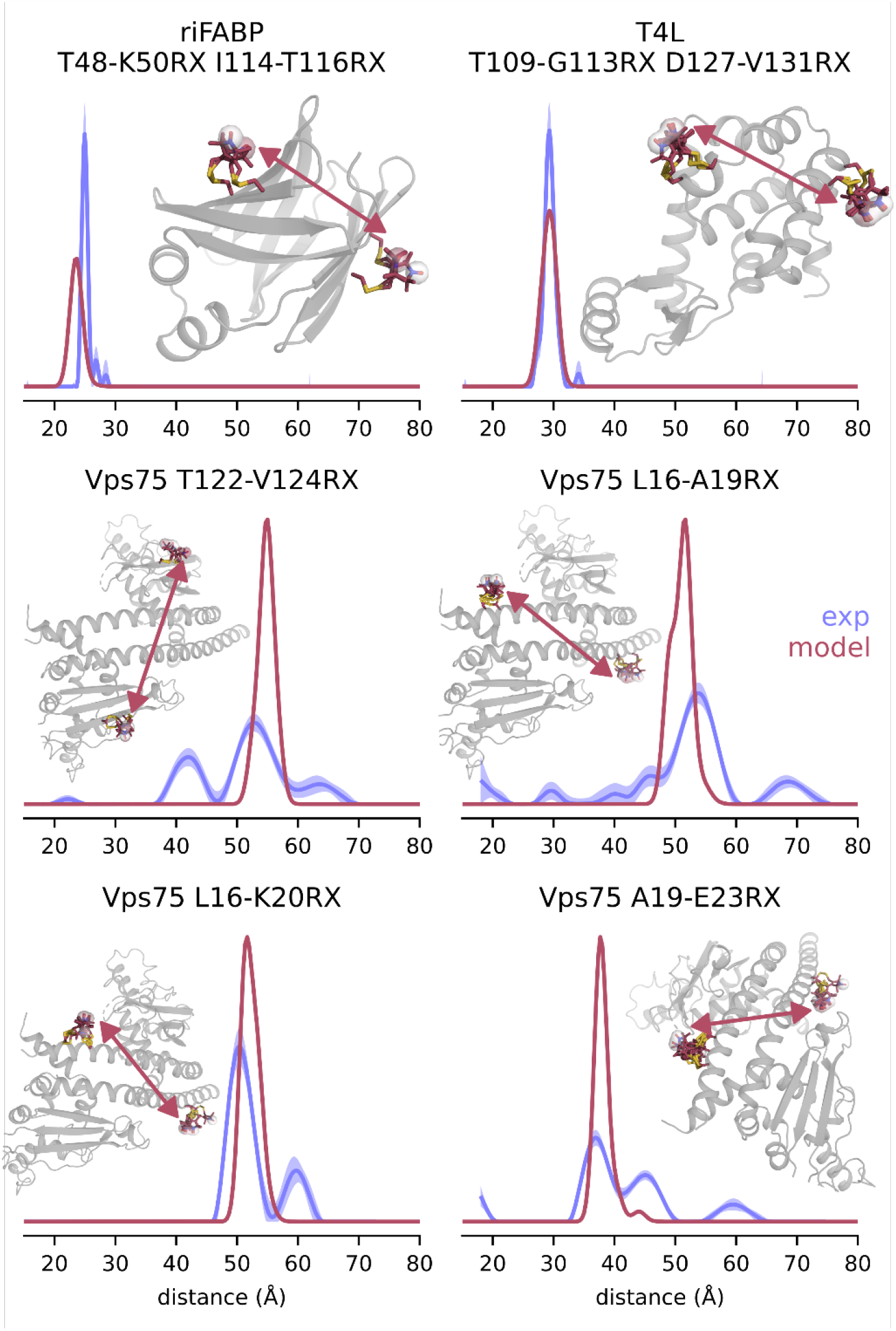
Bifunctional spin label modeling predictions of DEER distance distributions. Experimental distance distributions are shown in blue with 95% confidence intervals shown as semitransparent bands. Spin label model predictions are shown in red. Insets show the labeled protein (gray) with attached spin label ensembles shown as sticks with light blue carbons and nitrogen, sulfur, and oxygen are colored as in Fig. 1.

The results for site pairs in contexts that used the aggregate rotamer library are shown in Fig. 5 and Fig. S4. The predictions have a similar accuracy, with a MAD of 1.3 Å when comparing the modes of the distributions and a MAD of 2.3 Å when comparing the average distance. Again, the most pronounced differences resulted from additional modes in the experimental distributions that were not present in the predicted distributions. Despite these differences, the simulated distributions agree well with the primary modes of the experimental data, suggesting our approach is applicable in the absence of context-specific rotamer libraries.

**Fig. 5.**
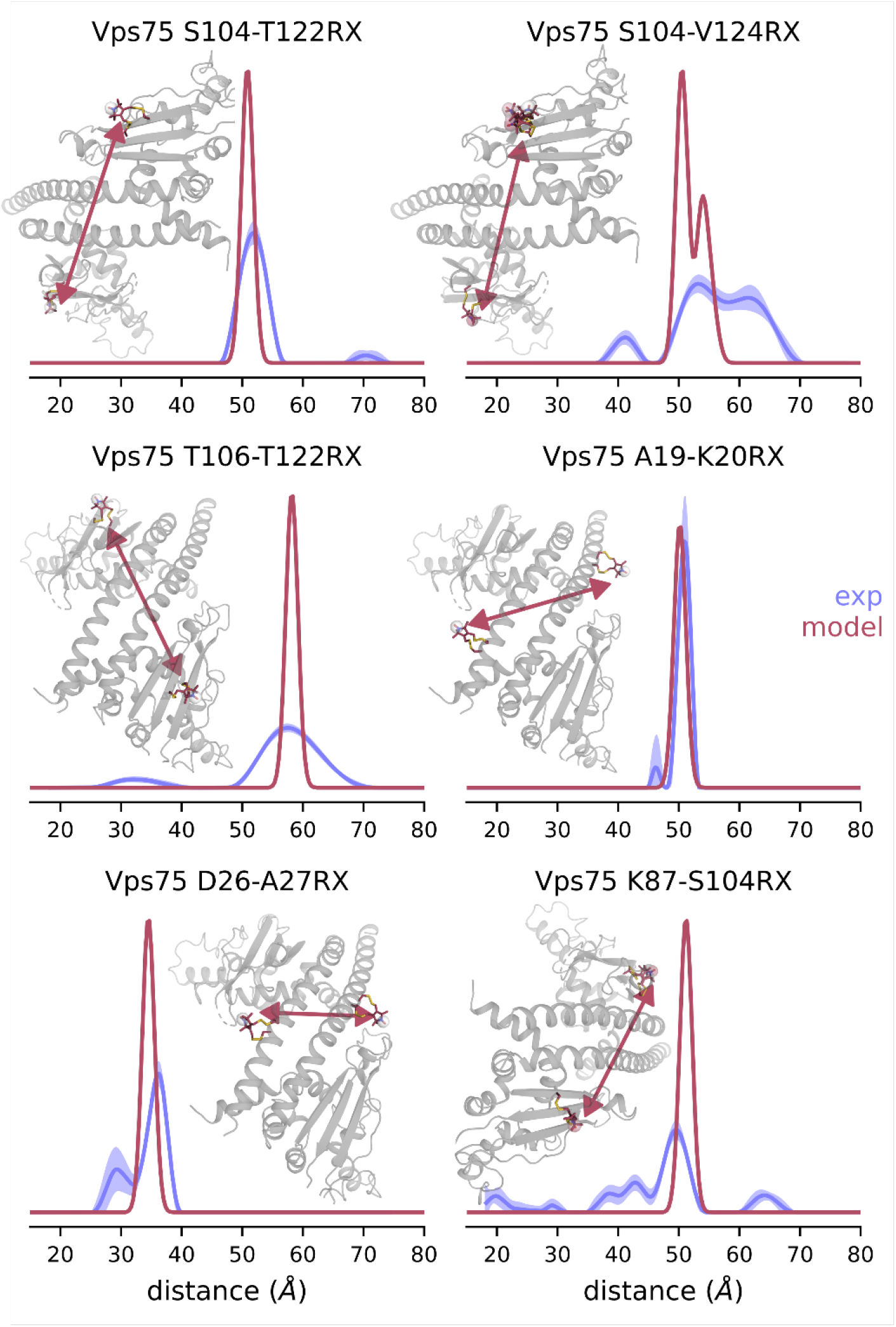
Bifunctional spin label modeling predictions of DEER distance distributions with the aggregate rotamer library. Experimental distance distributions are shown in blue with 95% confidence intervals shown as semitransparent bands. Spin label model predictions are shown in red. Insets show the labeled protein (gray) with attached spin label ensembles shown as sticks with light blue carbons and nitrogens, sulfurs and oxygens colored as in Fig. 1.

Overall, our approach appears to model the bifunctional RX label with similar accuracy to popular monofunctional label modeling methods [67]. While the Vps75 predictions differ slightly from the previously published MD simulations [56], they seem to recapitulate the general trends of the experimental data. It is important to note that we cannot make a direct comparison between our method and the previously reported MD simulations due to differences in the processing of experimental data.

Except for the T4L and riFABP data, all experimental distributions are considerably broader than the predicted distributions, suggesting there is additional backbone flexibility that is not captured by the Vps75 crystal structure. Accordingly, we expect that the bimodal distance distributions observed for the Vps75 constructs are also due to backbone flexibility, which would explain why we consistently were able to predict the primary mode of the distance distributions but not the secondary mode. As discussed above, it is unsurprising that this backbone flexibility is not captured by modeling spin labels on the crystal structure alone. We believe this caveat illustrates the potential of using more restrictive spin labels like RX to better resolve protein backbone flexibility while simultaneously using spin label and protein modeling to develop models of protein conformational landscapes. An alternative explanation is that a significant fraction of the bifunctional spin label was only attached at one site. In this case, a more accurate modeling approach may be to perform labeling with a mixed monofunctional-bifunctional rotamer library where the monofunctional component is composed of partially labeled spin label structures.

Taken together, the results shown in Figs. 4 and 5 suggest that our approach to modeling bifunctional spin labels can be used to predict protein SDSL EPR experimental results with reasonable accuracy.

### 3.2 Model Predictions of Crystal Structure

To further validate our modeling approach, we attempted to model RX on the α4 site pair T115-T119 of T4L and compare it with an existing crystal structure of the spin-labeled protein (PDB: 3L2X) [55]. Figure 6A shows the most probable (highest weighted) T115-T119RX model, generated using the 3L2X PDB file, in comparison to the experimental structure. The model side chain conformation comes quite close to the one from the crystal structure, with a 1.6 Å heavy atom (excluding β-carbons) RMSD. Approximating the spin center to be halfway between the nitrogen and the oxygen of the nitroxide moiety, places the modeled spin center only 1.5 Å from the crystal structure spin center. Notably, our modeling process produced an ensemble of 7 models, ranging from 1.3 to 1.6 Å heavy atom RMSD from the crystal model and 1.0 to 1.5 Å deviation of the approximate spin centers, suggesting consistent agreement between the ensemble and the crystal structure.

**Fig. 6.**
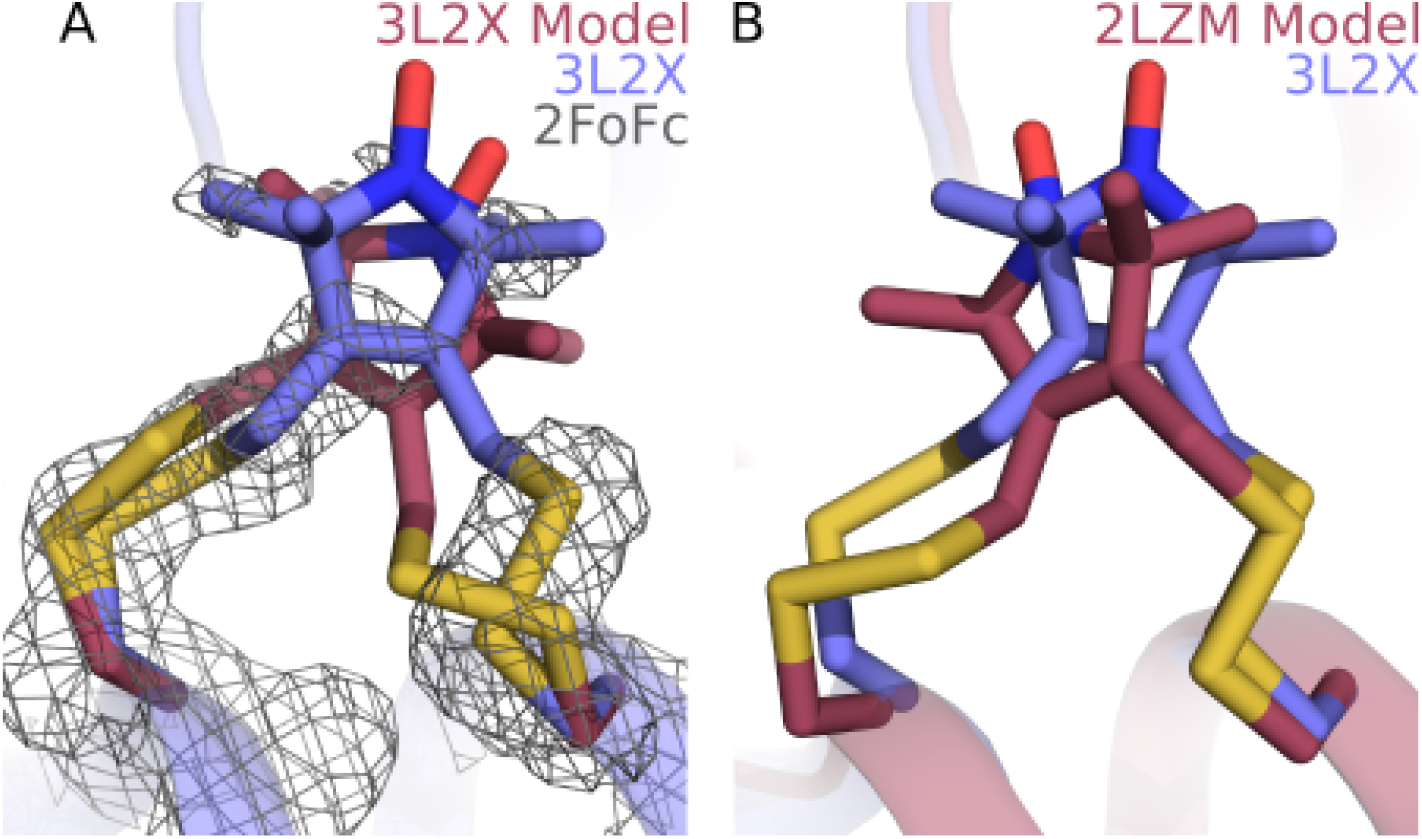
Comparison of T4L T115-T119RX spin label models (cartoons with dark red backbone and carbon atoms) with experimental crystal structure 3L2X (light blue backbone and carbon atoms). Nitrogen, sulfur and oxygen atoms are colored as in Fig. 1. A) Comparison of RX modeled using the 3L2X crystal structure. The 3L2X 2*F*_*o*_*–F*_*c*_ map is shown as a gray mesh contoured at 1.0 σ. B) Comparison of RX modeled using 2LZM crystal structure.

Notably, the 3L2X crystal structure differs slightly from the 2LZM crystal structure in the backbone and side chains near the labeling site. These small differences likely arise frequently during spin labeling experiments, and therefore, it is crucial that modeling methods are resilient to such perturbations and can generate consistent results despite the variations.

To evaluate the prediction power of our modeling method when using a native structure, we also modeled T115-T119RX on the 2LZM structure of unlabeled T4L and compared it to the RX label on the 3L2X structure (Fig. 6B). The most probable model came within 1.5 Å heavy-atom RMSD to the crystal structure and a 1.0 Å deviation of the approximated spin centers, despite the β-carbon of site 115 differing 1.2 Å between crystal structures. This model ensemble includes nine structures ranging from 1.5 to 2.7 Å heavy atom RMSD from the crystal model and 1.0 to 1.5 Å deviation of the approximate spin centers. These results suggest minimal influence of minor perturbations to the protein backbone and neighboring side chains on RX side chain modeling.

Cumulatively, the comparisons with existing crystal structures further illustrate the validity of our modeling method.

## 4. Conclusions

The restriction of side chain conformational dynamics offered by bifunctional labels like RX can result in significantly improved resolution of protein backbone dynamics. Full quantitative utilization of this improved backbone resolution requires accurate and efficient methods to model the spin label ensemble on multiple scales. We have developed a novel approach for modeling bifunctional spin label ensembles like RX and have shown that it has similar experimental prediction accuracy to existing monofunctional spin label modeling methods. While methods based on MD and metadynamics can also be used to model bifunctional labels [56,68,72], our approach is considerably faster (see Methods), is readily integrable with 3^rd^ party modeling software [84], and may be more amenable to rapid site pair screening to aid experimental design. Furthermore, our approach can be used to simplify more advanced methods by generating geometrically valid precursor models often required for other modeling methods [56,68].

Future advancements in bifunctional spin label modeling will seek to expand this method to other bifunctional spin labels [57,58,93] and to bifunctional chromophores for fluorescence resonance energy transfer (FRET) [94].

## Author Contributions

The modeling methodology and experimental design were developed by MHT and SS. The software was implemented by MHT. The original draft was prepared by MHT and edited by MHT and SS.

## Acknowledgments

The research presented here was supported by the National Institute of Health grant R01 GM125753 (SS). T4L and riFABP DEER data were recorded by the Hubbell group and generously provided by Michael Bridges. The Vps75 DEER data were recorded by the groups of David G. Norman and Graham M. Smith and generously provided by Hassane El Mkami. High-performance computing was performed through the Hyak computing cluster at the University of Washington.

