## Supplemental data for "A novel approach to modeling side chain ensembles of the bifunctional spin label RX"

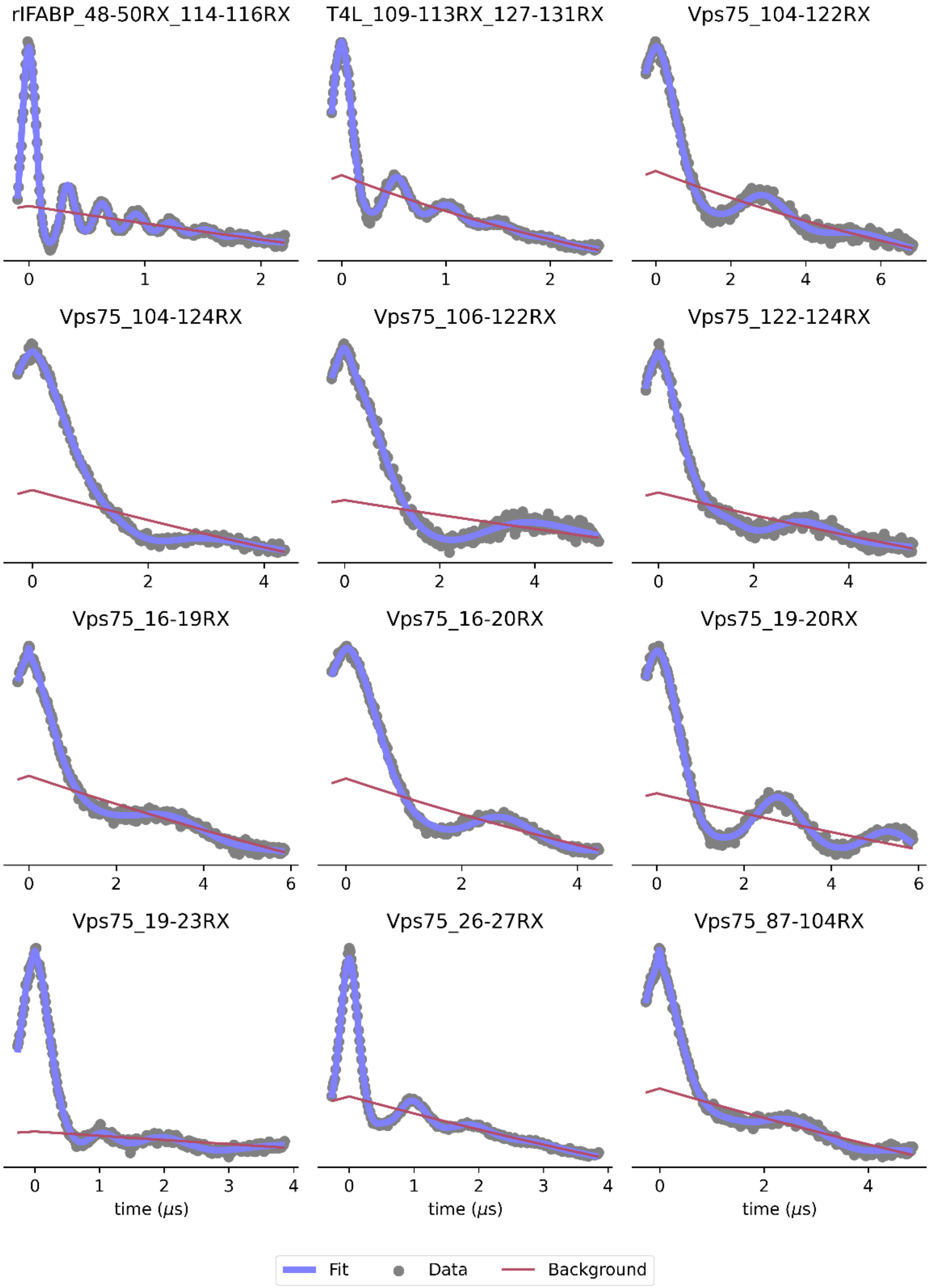


**Figure S1**. Raw data, foreground, and background fits.

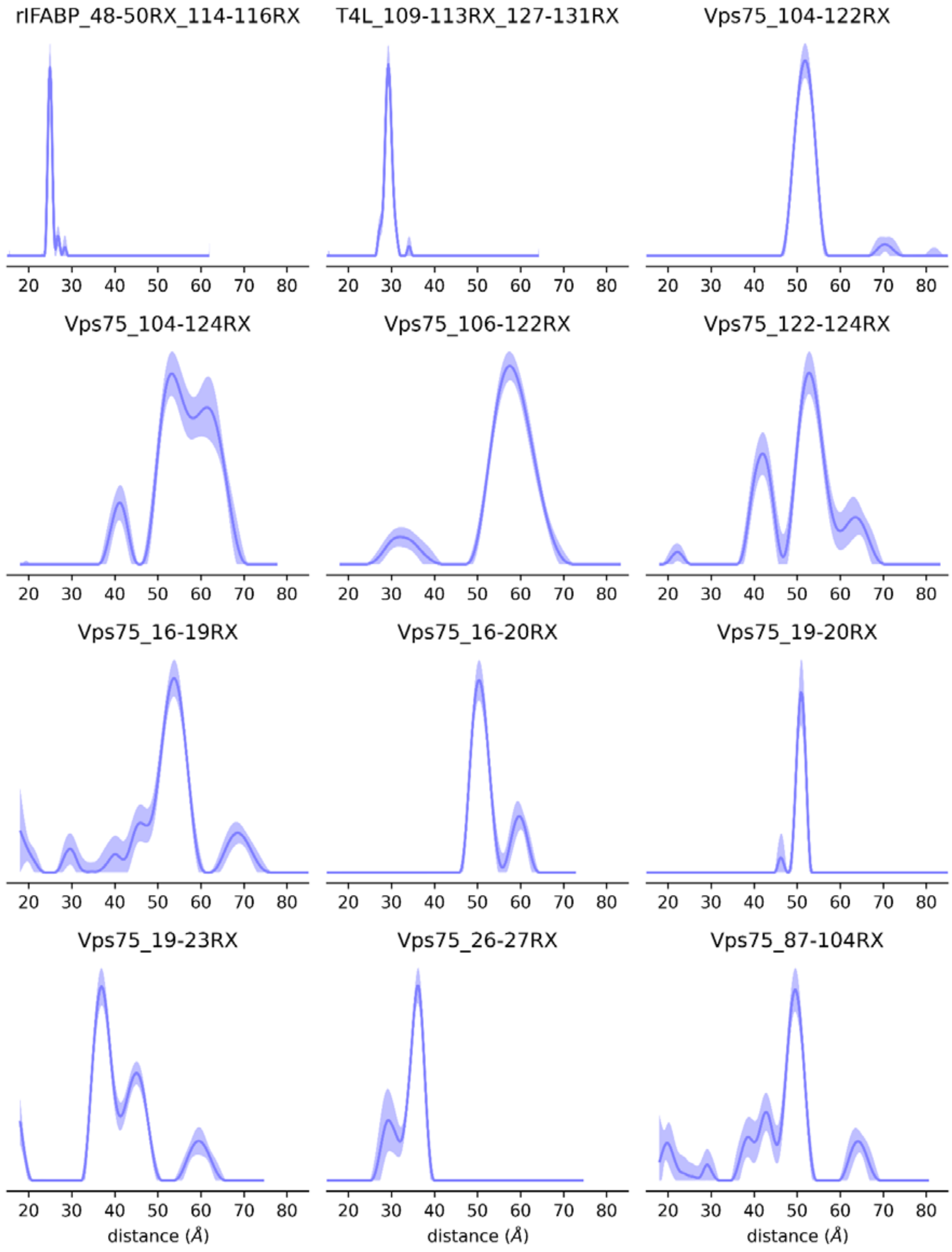


**Figure S2**. Experimental distance distributions, including 95% confidence interval bands.

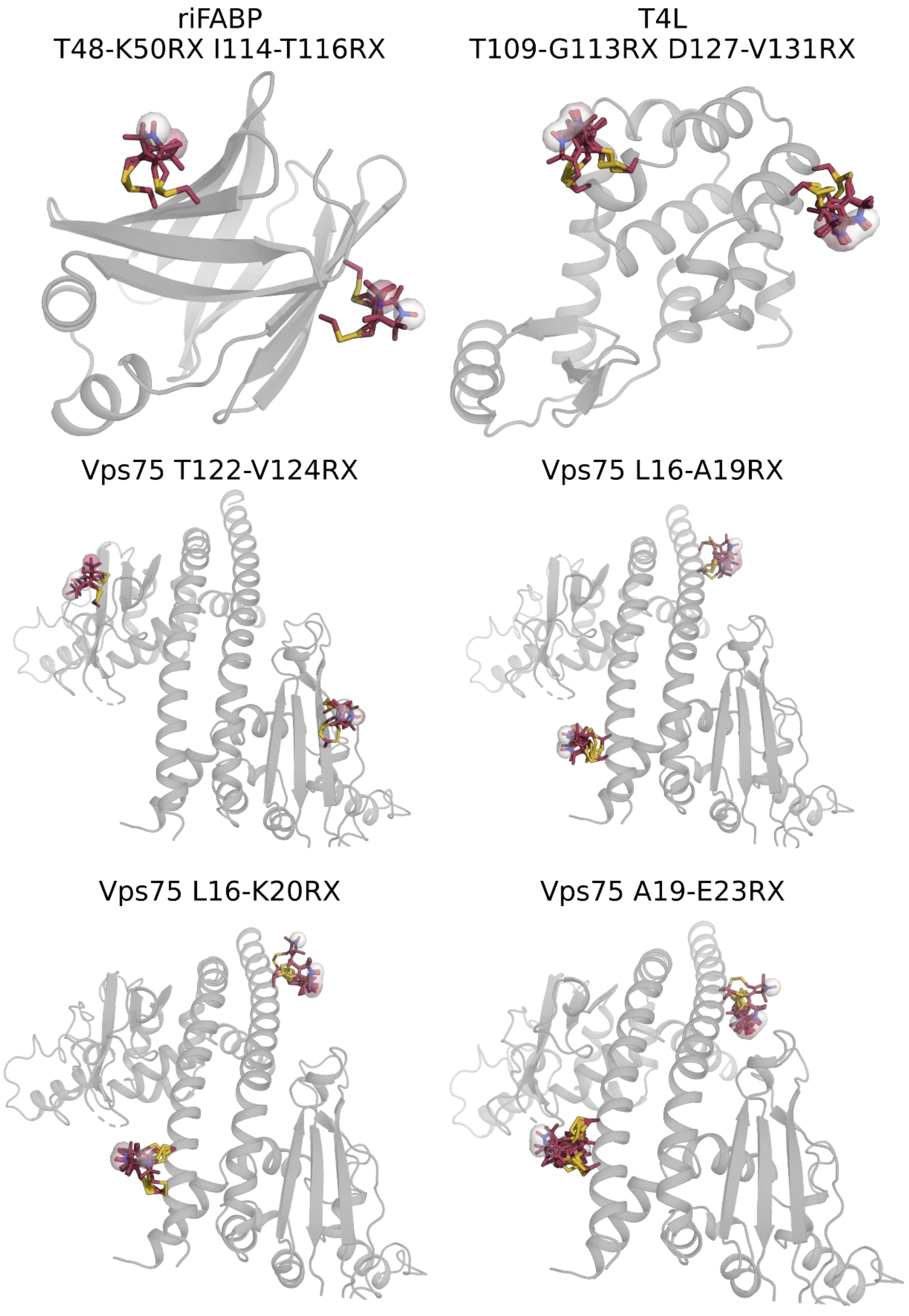


**Figure S3**. Bifunctional spin label ensembles attached to PDB structures. PDB structures of riFABP (2IFB), T4L (2LZM), and Vps75 (2ZD7) are shown as gray cartoons. RX Label ensembles are shown as sticks with red carbons.

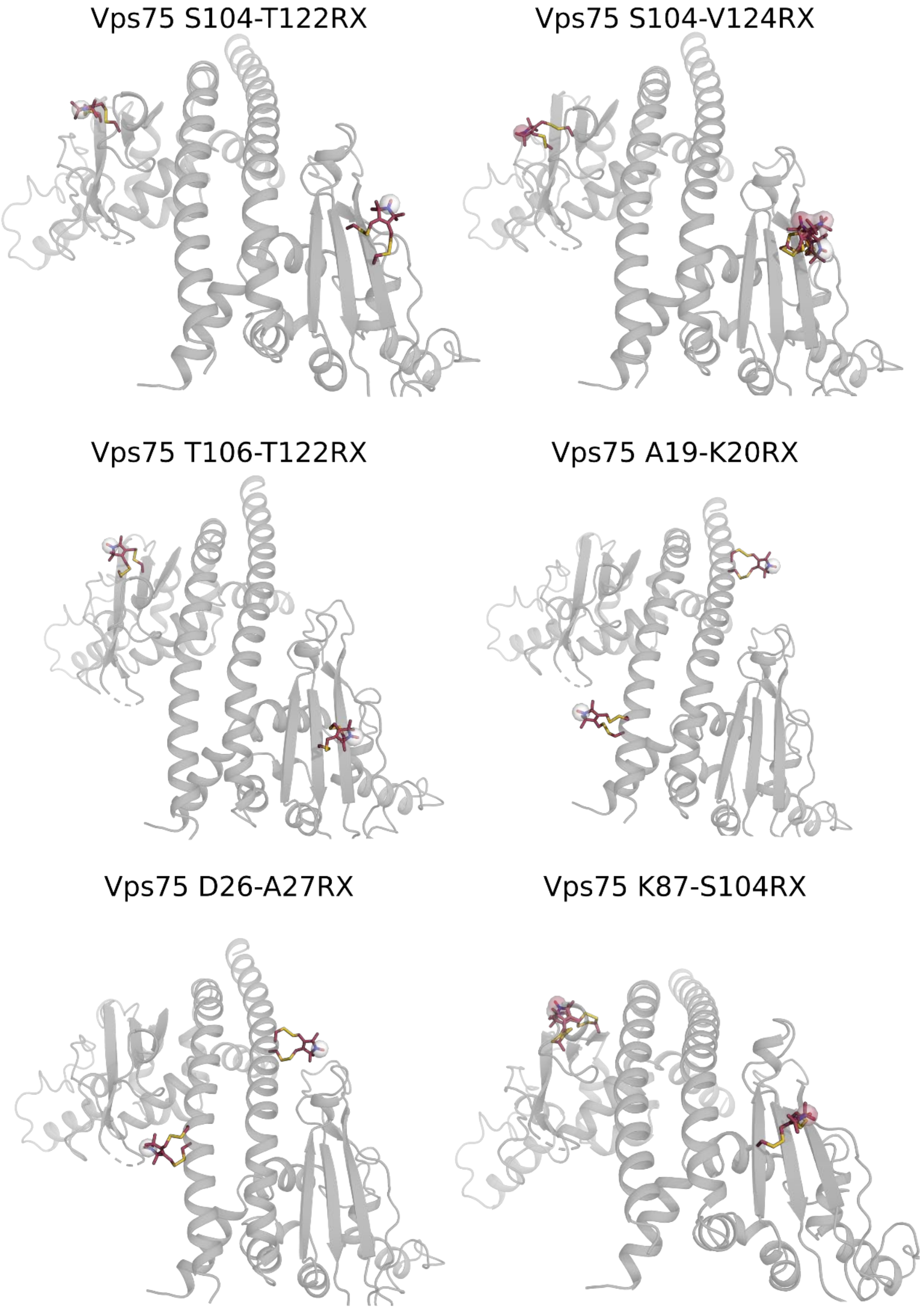


**Figure S4**. Bifunctional spin label ensembles attached to PDB structures. PDB structures of Vps75 (2ZD7) are shown as gray cartoons. RX Label ensembles are shown as sticks with red carbons.

**Table S1**. Experimental and fit parameters. Raw data provided by Stevens et al. and Fleissner et al.

| Sample | $\Lambda$[a] | Scans | $\Delta t$[b] (ns) | SNR[c] | $t_0$ offset (ns) | $\tau_2$ (μs) | SRT[d] (ms) | $A$[e] | $B$[f] |
|---|---|---|---|---|---|---|---|---|---|
| rIFABP_48-50RX_114-116RX | 0.39 | 484 | 8 | 81 | 98.4 | 2.248 | 1.02 | 0.02 | 0.46 |
| T4L_109-113RX_127-131RX | 0.45 | 1145 | 8 | 101 | 96 | 2.5 | 1.02 | 0.04 | 0.25 |
| Vps75_104-122RX | 0.40 | 63 | 20 | 37 | 266 | 7 | 4.08 | 0.45 | 0.06 |
| Vps75_104-124RX | 0.30 | 121 | 23 | 49 | 254.4 | 4.5 | 4.08 | 0.69 | 0.04 |
| Vps75_106-122RX | 0.37 | 89 | 20 | 32 | 276 | 5.5 | 4.08 | 2.72 | 0.05 |
| Vps75_122-124RX | 0.35 | 107 | 20 | 33 | 270 | 5.5 | 4.08 | 0.81 | 0.05 |
| Vps75_16-19RX | 0.26 | 77 | 20 | 40 | 262 | 6 | 4.08 | 0.45 | 0.02 |
| Vps75_16-20RX | 0.35 | 5 | 20 | 63 | 250 | 4.5 | 4.08 | 0.29 | 0.06 |
| Vps75_19-20RX | 0.32 | 76 | 20 | 39 | 254 | 6 | 4.08 | 0.04 | 0.66 |
| Vps75_19-23RX | 0.44 | 73 | 16 | 74 | 267.2 | 4 | 4.08 | 0.45 | 0.03 |
| Vps75_26-27RX | 0.32 | 20 | 16 | 121 | 273.6 | 4 | 4.08 | 0.14 | 0.37 |
| Vps75_87-104RX | 0.31 | 104 | 20 | 52 | 270 | 5 | 4.08 | 0.24 | 0.02 |

[a] – Modulation depth
[b] – Timestep
[c] – Signal to Noise Ratio (estimated)
[d] – Shot repetition time
[e] – Smoothing regularization parameter
[f] – Compactness regularization parameter
